# Multiple Cross Displacement Amplification Coupled with Gold Nanoparticles-Based Lateral Flow Biosensor for Detection of the Mobilized Colistin Resistance Gene *mcr-1*

**DOI:** 10.1101/579904

**Authors:** Lin Gong, Ernan Liu, Jie Che, Juan Li, Xiaoli Liu, Huiqiong Xu, Jiansheng Liang

## Abstract

Fast dissemination of the mobilized colistin resistance (*mcr*) gene *mcr-1* in *Enterobacteriaceae* causes a huge threat to the treatment of severe infection. In the current report, a multiple cross displacement amplification (MCDA) coupled with the detection of amplified products by gold nanoparticles-based lateral flow biosensor (LFB) assay (MCDA-LFB) was established to identify the *mcr-1* gene with simpleness, rapidity, specificity and sensitivity. The MCDA-LFB assay was performed at a isothermal temperature (63°C) for only 30 min during the amplification stage, and the reaction products were directly identified by using LFB which obtained the result with 2 min. The entire process of experiments, from templates extraction to result judging, was accomplished less than 60 min. For the analytical specificity of this method, all of the 16 *mcr-1*-producing strains were positive, and all of the non-*mcr-1* isolates got the negative results. The sensitivity of *mcr-1*-MCDA-LFB assay was as little as 600 fg of plasmid DNA per reaction in pure culture, and approximately 4.5×10^3^ CFU/mL (~4.5 CFU/reaction) in fecal samples spiked with 100 μl of strains. Therefore, this technique established in the present study is suitable for the surveillance of *mcr-1* gene in clinic and livestock industry.

## Introduction

The rapid increase of carbapenem-resistant Enterobacteriaceae (CRE) expressing *Klebsiella pneumoniae* carbapenemase(KPC), New Delhi metallo-blactamase(NDM) and oxacillinase(OXA) OXA-48 has risen serious concerns in clinic. Colistin, a ‘last resort’ antibiotic, has a crucial role for treating the infection caused by those species (1). Therefore, that the number of colistin consumptions are increasing along with the global augment of CRE bring about the risk of emerging resistance (2).

Resistance to colistin was linked with chromosomal resistance mechanisms in varieties of strains in the past (3). Since a new mobilized colistin resistance gene, mcr-1, carried by plasmid in an Escherichia coli was first reported in China in 2015 (4), which has been identified in numerous countries. China, Germany and Vietnam carry an important proportion of positive samples (5). The *mcr-1*-positive bacterial species include *Salmonella enterica, Escherichia coli, Escherichia fergusonii, Enterobacter aerogenes, Klebsiella pneumoniae, Citrobacter braaki* and *Klebsiella aerogenes* (5-11). Besides discoveried in Clinical samples, *mcr-1* is also detected from environmental settings: meat and vegetable products purchased from markets, Animal feces collected from farms, fecal samples of pets gathered from pet hospital, river water and seawater (12). The wide dissemination of *mcr-1* across diversified species is benefited from many types of *mcr-1*-bearing plasmids covering IncHI2, IncI2, IncFI, IncX4 and IncX1-X2 hybrid type (13-17). Similarly, the genetic environments of *mcr-1* gene also impact its transmission. A global data set of roughly 500 isolates producing *mcr-1* analysed by whole-genome sequencing (WGS) has revealed that an initial mobilized event of *mcr-1* is mediated by an ISApl1-*mcr-1-orf*-ISApl1 transposon around 2006 (5). The horizontal transfer of *mcr-1* gene causing inflation of colistin-resistant isolates will lead to the shortage of effective measures for treating infections with multidrug-resistant bacteria. Therefore, a rapid, sensitive and specific diagnostic assay for *mcr-1* detection is imperative to devise.

Currently, sereval categories of molecular diagnostic methodologies including conventional PCR and real-time PCR methods have been devised to identify *mcr-1* gene (18). Nevertheless, the requirements of highly sophisticated devices, Strictly experimental environments and well-trained professionals restrict those techniques to apply in resource-challenged areas and “on-site” detection (19). Recently, multiple cross displacement amplification (MCDA), a novel nucleic acid amplification technique, has been utilized in detection of bacterial agents, such as *Listeria monocytogenes* and methicillin-resistant *Staphylococcus aureus* (MRSA) (20, 21). With the advantages of rapidity, specificity and sensitivity, MCDA operated in a simple heater can yield amplifcons from a few colonies (20, 22), the amplicons are identified by a gold nanoparticles-based lateral flow biosensor (LFB) subsequently.

In this study, a MCDA-LFB assay for the rapid detection of *mcr-1* was established, and the sensitivity and specificity of above method in pure culture and in spiked fecal samples were analyzed.

## Materials and Methods

### Reagents and instruments

Bacterial genomes extraction kits were obtained from Beijing ComWin Biotech Co.,Ltd.(Beijing, China). QIAGEN plasmid kits and QIAamp fast DNA stool mini kits were purchased from Qiagen Co., Ltd.(Beijing, China). Isothermal amplification kits (including reaction buffer and Bst DNA polymerase 2.0), colorimetric indicator, and disposable lateral flow biosensor were provided by BeiJing-HaiTaiZhengYuan Technology Co., Ltd.(Beijing, China). The heating thermostat (MTH-100) was purchased from Hangzhou MiU Instruments Co., Ltd.(Hangzhou, China). The UV transilluminator (UVsolo touch) was obtained from Analytik Jena(Jena, Germany). Nanodrop instrument (ND-2000) was purchased from Thermo Fisher Scientific Co., Ltd. (Massachusetts, America).

### Isolates and genomic template preparation

A total of 51 organisms consisting of 16 *mcr-1*-postive isolates and 35 non-*mcr-1* bacteria were used in the this study (**Table 1**). The *mcr-1*-postive bacterial species included 5 *Escherichia fergusonii,* 10 *Escherichia coli* and 1 *Salmonella enteritidis.* The mcr-1-negative isolates involved KPC-2-postive stains(*Klebsiella pneumoniae* and *Pseudomonas aeruginosa*), NDM-1-postive isolates(*Escherichia coli, Enterobacter cloacae* and *Klebsiella pneumoniae*), NDM-5-producing *Escherichia coli,* IMP-4-producing *Escherichia coli,* and *mcr-1/*carbapenemase-negative species (*Acinetobacter baumannii, Pseudomonas aeruginosa, Serratia marcescens* and *Escherichia coli*). According to the handling instruction, the genomes DNA of all strains was extracted by bacterial genomes extraction kits, the plasmid DNA of *mcr-1-*producing *Escherichia fergusonii* (ICDC-ZG2016M34-3) which was acted as a representative sample for optimization of reaction condition and sensitivity detection was acquired by QIAGEN Plasmid Kits, and quantified by a Nanodrop ND-2000 instrument.

**TABLE 1.**
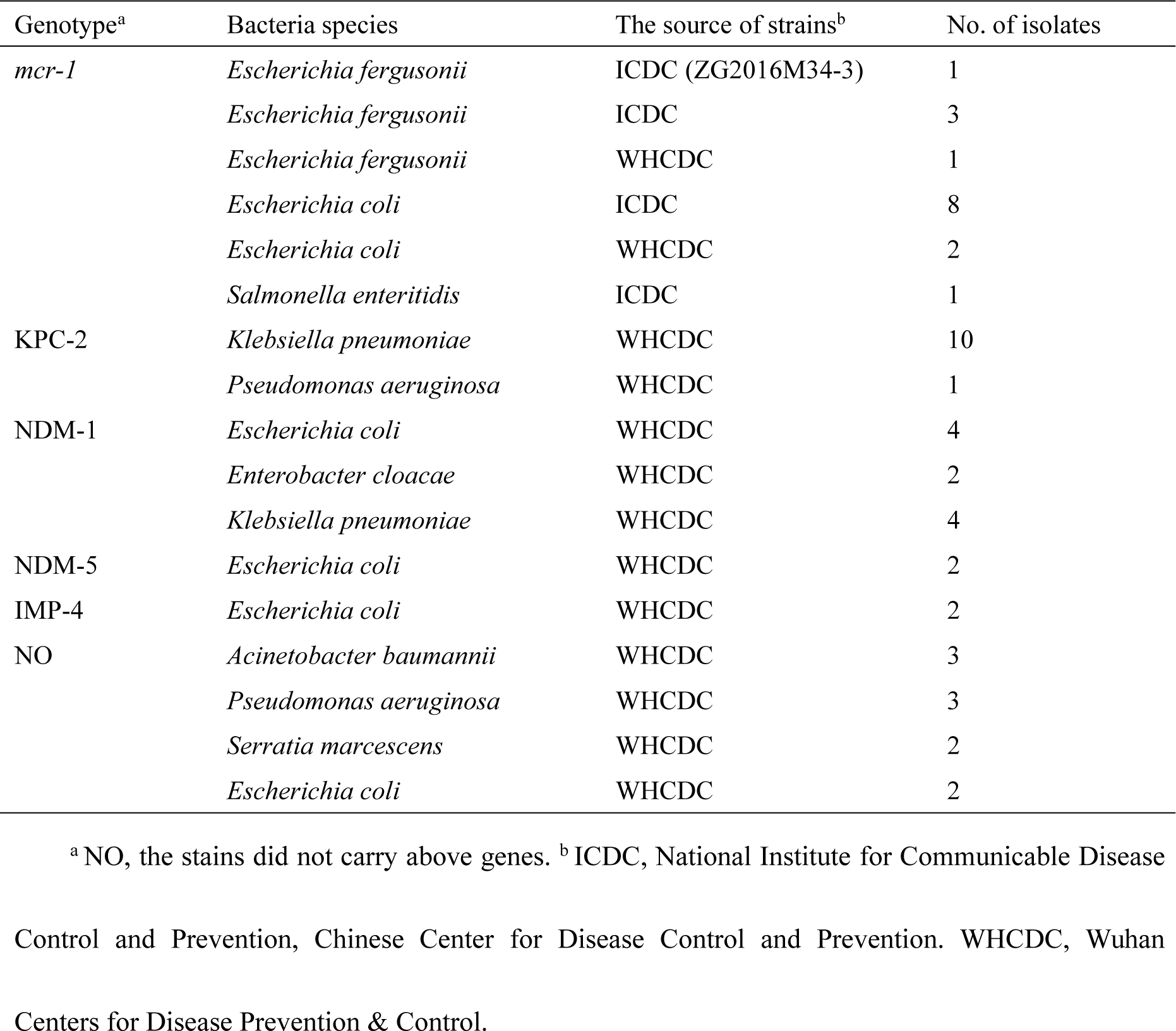
Strains used in this study.

### Primers design of MCDA assay

Two softwares named Primer Primer 6.0 and PrimerExplorer V4 were used to design the five pairs of MCDA primers based on *mcr-1* gene. The dimer and hairpin structures of all primers were detected by Integrated DNA Technologies design tools (23), and the specificity of which was analyzed by using Basic Local Alignment Search Tool (Blast). The relevant information of primers pairs (F1, F2, C1, C2, D1, D2, R1, R2, CP1 and CP2) about positions and sequences is displayed in **Fig. 1** and **Table 2**. Furthermore, The FITC (Fluorescein isothiocyanate) and biotin labeled at 5’end of the C1 and D1 primers, respectively, and the new primers were named as C1* and D1*. All of the primers were synthesized and purified by Sangon Biotechnology Co., Ltd.(Shanghai, China) at HPLC purification grade.

**TABLE 2.**
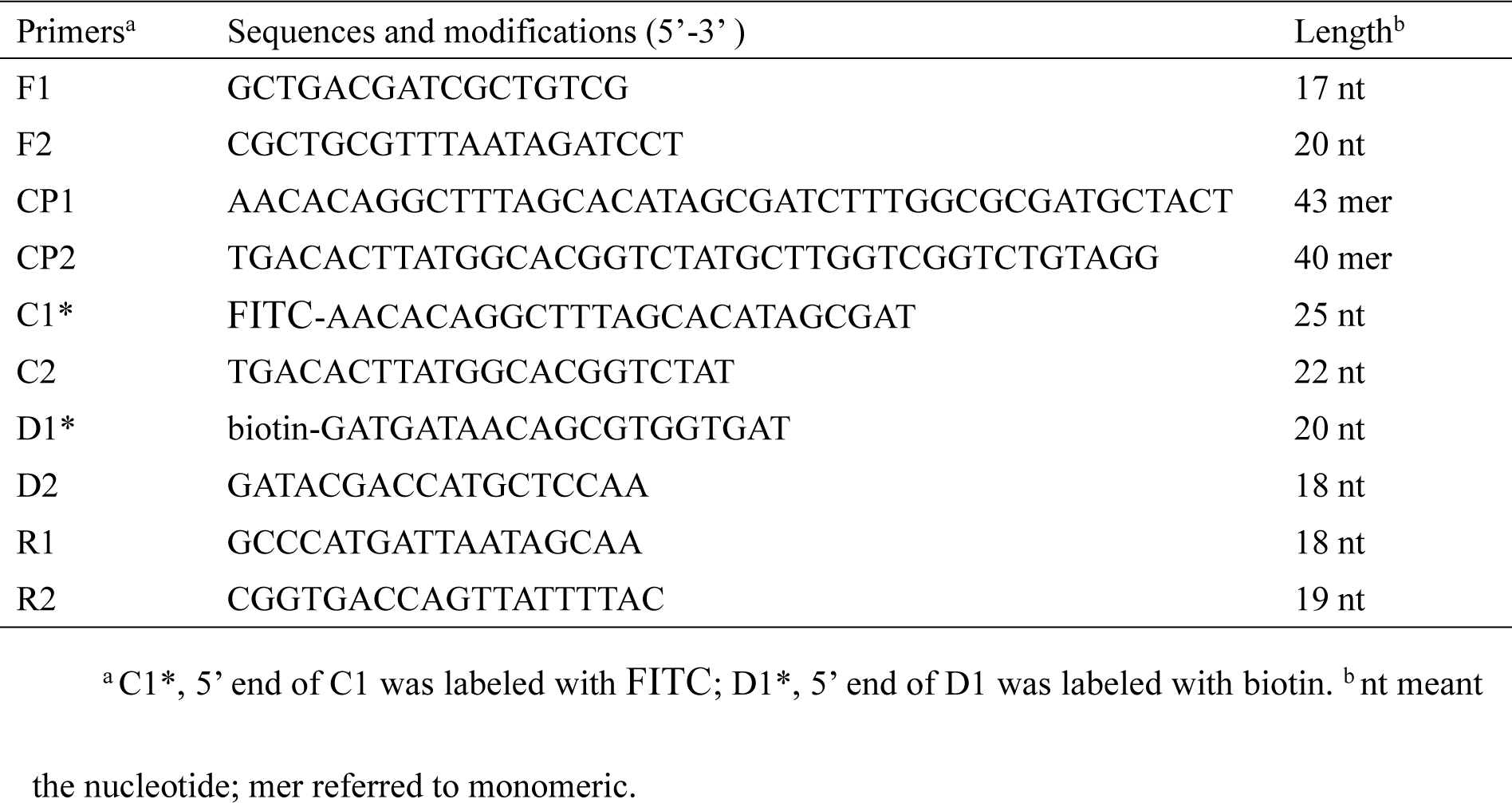
Primers of the MCDA assay to identify the *mcr-1* gene.

**FIG 1.**
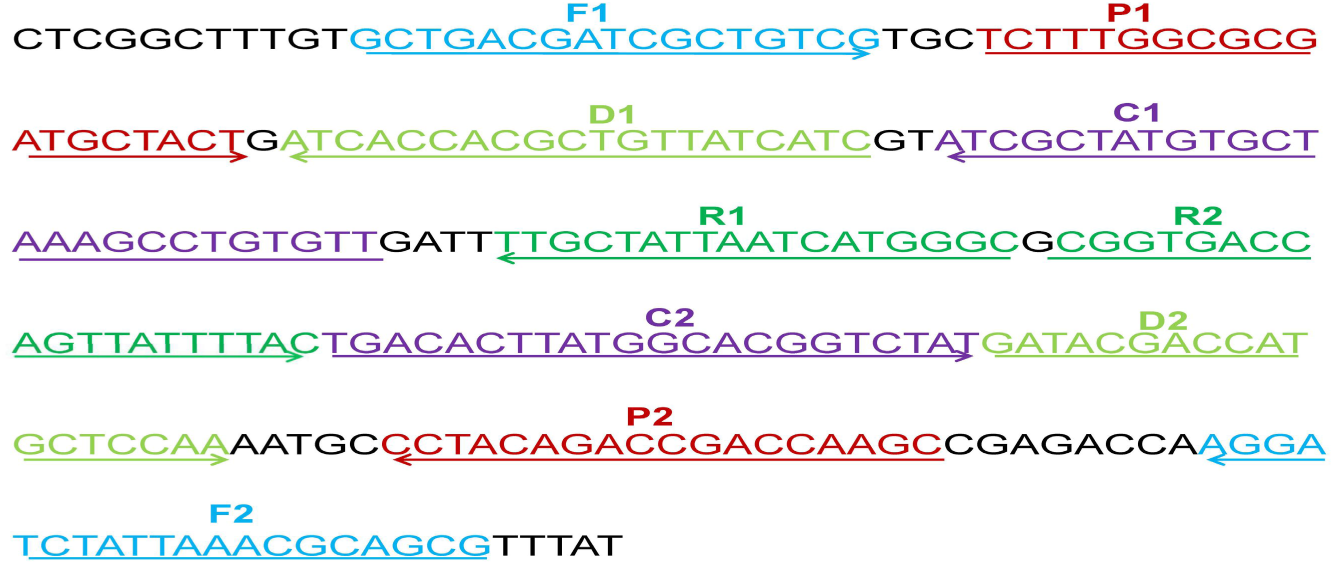
The location of the five primers pairs on the partial *mcr-1* gene sequence. Different pairs of primers were represented by various of colors. The arrowed lines showed the amplification direction of primers.

### The standard MCDA assay

The MCDA reaction systems were performed according to the previous studies (23). Each reaction, the total volumes of 25 μL, included reaction buffer (12.5 μL), Bst DNA polymerase 2.0 (1 μL), colorimetric indicator (1 μL), cross primers CP1 and CP2 (1.6 μM each), displacement primers F1 and F2 (0.4 μM each), amplification primers C1, C2, D1, D2, R1 and R2 (0.4 μM each), and 1 μL of DNA template. Mixtures including 1 μL of DNA of NDM-1-positive *Escherichia coli* (WHCDC-WH67) and KPC-2-producing *Klebsiella pneumoniae* (WHCDC-WH108) were regarded as the negative controls, and 1μL of distilled water contained in mixtures was served as the blank control. To assess the optimal reaction temperature of *mcr-1*-MCDA primers, the MCDA amplification systems were executed with a constant temperature in the range of 60 °C to 67°C for 40 min.

The MCDA reaction products were analyzed by three detection methods containing 2% agarose gel electrophoresis, colorimetric indicator, and LFB (20). When employing gel electrophoresis, 3 μL of reaction mixtures were run at 110 volts for 60 min. A ladder of multiple bands could be observed in the positive reactions, but not in the negative and blank controls. Reaction products were detected by using colorimetric indicator, the color of amplified products remained unchanged. Nevertheless, the negative and blank controls reactions changed from blue to colorless. The material, theory and operation procedure of LFB were previously described by Wang et al. (20). 0.2 μL of amplicons followed by three drops of the running buffer consisting of 1% Tween 20 and 0.01mol/L phosphate-buffered saline were added to the well of sample pad (19). After one to two minutes, two red lines named test line (TL) and control line (CL) respectively, could be visualized in positive products, while only the CL was observed for the negative and blank control.

### Sensitivity and specificity of the *mcr-1-*MCDA-LFB assay

The plasmid DNA of *Escherichia fergusonii* ICDC-ZG2016M34-3 was serially diluted (6ng, 60pg, 600fg, 60fg, and 6fg per μl) for sensitivity analysis of *mcr-1*-MCDA-LFB detection. The colorimetric indicator and agarose gel electrophoresis were carried out simultaneously. Each test was repeated three times. The specificity of *mcr-1*-MCDA-LFB assay was evaluated with the DNA templates of 16 *mcr-1-*producing strains and 35 non-*mcr-1* strains (**Table 1**), The specificity evaluations were confirmed twice.

### The optimal amplification time

In order to screen the optimal time for the *mcr-1*-MCDA-LFB assay, The MCDA mixture was completed at the reaction temperature in the range from 10 min to 40 min at 10 min intervals. Subsequently, the MCDA products were detected by LFB detection. Each amplification time was operated at two times.

### *Mcr-1*-MCDA-LFB detection in fecal samples

The fecal samples were obtained from a healthy man in Wuhan, China. *Mcr-1* gene was not detected in those samples accroding to the microbial culture and PCR identification. The volumes of 100 μL were taken out from the *mcr-1*-positive *Escherichia fergusonii* ICDC-ZG2016M34-3 cultures when the optical density (OD) of that reached to 0.6. The suspensions were serially diluted (10^−1^ to 10^−8^), and the aliquots of 100 μL dilutions (10^−3^-10^−6^) were incubated on nutrient agar plates with three replicates, colony forming units (CFUs) were counted subsequently. 100 μl of diluted *mcr-1*-producing cultures (10^−2^ to 10^−7^) with known amounts (4.5 × 10^6^ to 4.5 × 10^1^ CFU/ mL) was respectively added to 0.2g of fecal sample and mixed well, DNA templates were extracted with the manufacturer’s protocol by using QIAamp fast DNA stool mini kits. The extracted genomic DNA was dissolved in 100μl of elution buffer, and 1 μl of which was used for MCDA-LFB detection as templates. A non-spiked feces sample was tested as negative control. The products of MCDA were also detected by colorimetric indicator and 2% agarose gel electrophoresis. The evaluation assay for limit of detection in fecal samples was conducted triplicate.

## Result

### The verification of *mcr-1* MCDA Products

The *mcr-1* MCDA assays were performed at a constant temperature (63°C) for 40 min to verify the availability of MCDA primers. Positive reaction appeared with DNA from *mcr-1*-producing *Escherichia fergusonii* (ICDC-ZG2016M34-3), but not with NDM-1-positive *Escherichia coli* (WHCDC-WH67), KPC-2-producing *Klebsiella pneumoniae* (WHCDC-WH108) and the blank contol (**Fig. 2**). Therefore, the primers of *mcr-1*-MCDA was suitable for establishment of the MCDA-LFB assay to detect *mcr-1* gene.

**FIG 2.**
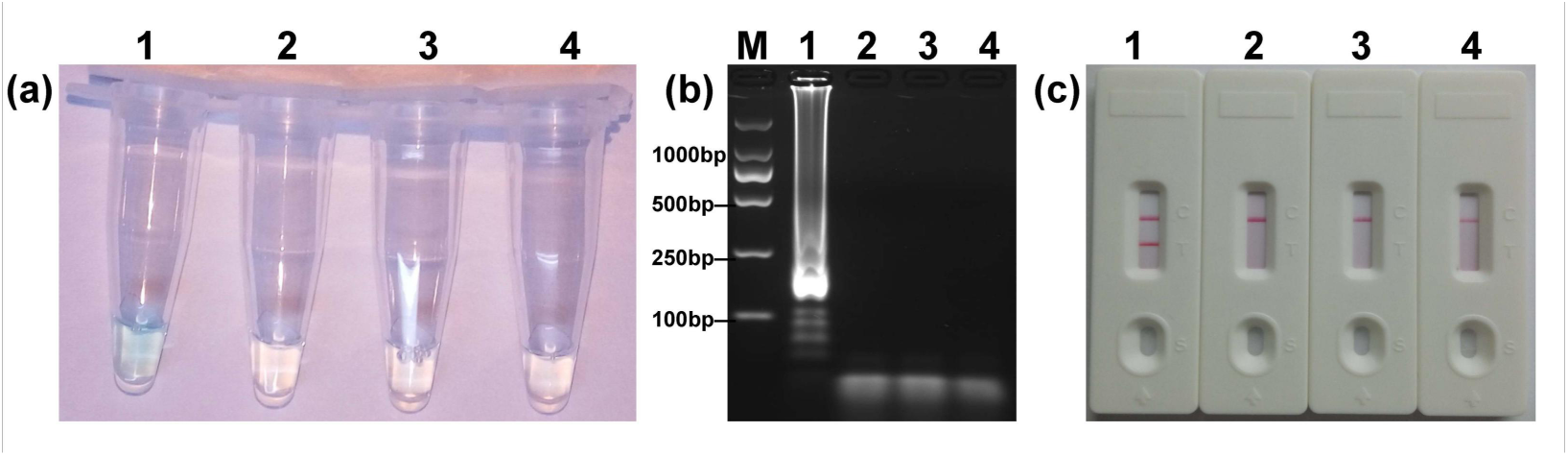
Confirmation of the *mcr-1*-MCDA-LFB assay. There were three detction methods to identify MCDA products: (a) colorimetric indicators, (b) agarose gel electrophoresis, (c) LFB. Four samples were tested: 1. *mcr-1*-positive strain of *Escherichia fergusonii* (ICDC-ZG2016M34-3); 2. NDM-1-positive *Escherichia coli* (WHCDC-WH67); 3. KPC-2-producing *Klebsiella pneumoniae* (WHCDC-WH108); 4. distilled water. Only the amplication with *Escherichia fergusonii* (ICDC-ZG2016M34-3) showed the positive.

### Temperature optimization for *mcr-1*-MCDA-LFB assay

To optimize the reaction temperature of MCDA-LFB assay during the amplification stage, the plasmid DNA of *Escherichia fergusonii* (ICDC-ZG2016M34-3) at the level of 6pg per reaction was used as the templates. A series of temperatures (ranging from 60°C to 67°C, with 1°C intervals) was compared for amplifying efficiency of MCDA-LFB assay by employing 2% agarose gel electrophoresis. The result showed that 62°C and 63°C were the better candidates for this method (**Fig. 3**). Therefore, the reaction temperature of 63°C was performed for the subsequent MCDA-LFB experiments.

**FIG 3.**
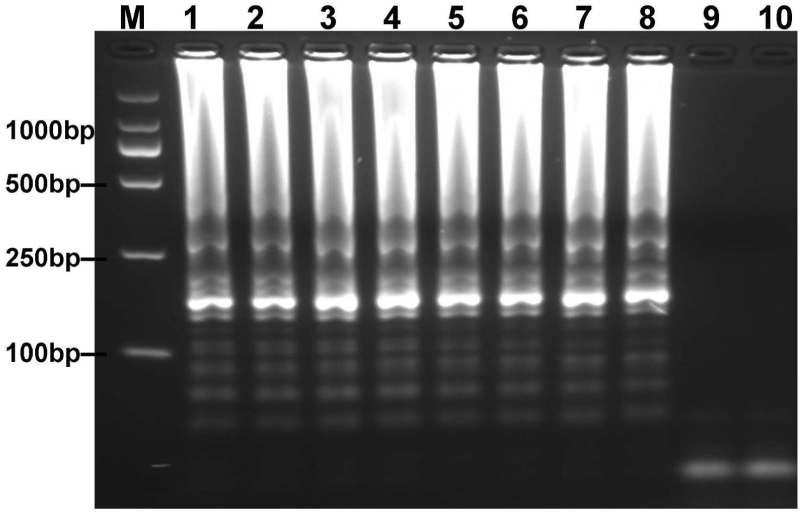
Optimal reaction temperature for *mcr-1*-MCDA assay was determined by using agarose gel electrophoresis. The plasmid DNA of *Escherichia fergusonii* (ICDC-ZG2016M34-3) (6pg) was amplified in different temperatures. Lane 1–8 represented the ampication temperature in the range from 60°C to 67°C (1°C intervals); lane 9, negative control (6 pg of *Escherichia coli* WHCDC-WH67 genomic DNA); lane 10, blank control (DW).

### Sensitivity and specificity of MCDA-LFB assay for *mcr-1*

To acquire the detection limit of this assay, serial dilutions of *Escherichia fergusonii* ICDC-ZG2016M34-3 plasmid DNA were used in *mcr-1*-MCDA-LFB assay. It indicated that the threshold was as little as 600 fg of plasmid DNA (**Fig. 4**). The same results were observed by using colorimetric indicator and agarose gel electrophoresis analysis.

**FIG 4.**
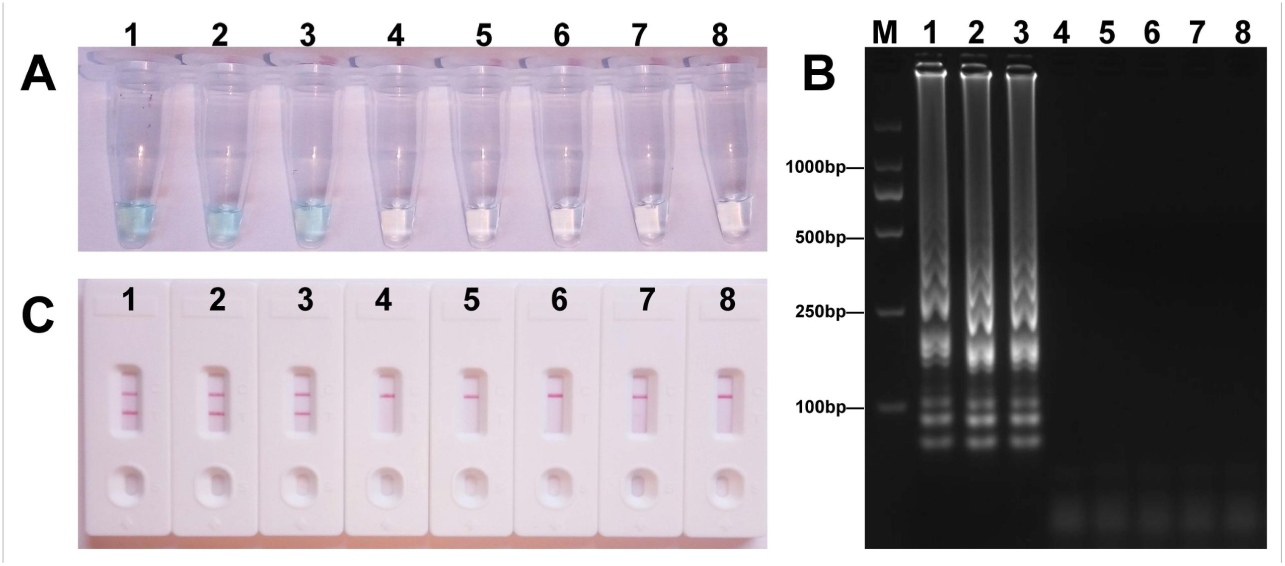
Sensitivity of the *mcr-1*-MCDA assay using serially diluted plasmid DNA with *Escherichia fergusonii* ICDC-ZG2016M34-3. Three detection techniques, including colorimetric indicators (A), gel electrophoresis (B) and LFB (C), were applied for testing MCDA products. Tubes (A)/lanes (B)/biosensors(C) 1-8 represented the plasmid DNA levels of 6 ng, 60 pg, 600 fg, 60 fg and 6 fg per reaction, negative control (NDM-1-positive *Escherichia coli* WHCDC-WH67 genomic DNA, 6 pg per reaction), negative control (KPC-2-producing *Klebsiella pneumoniae* WHCDC-WH108 genomic DNA, 6 pg per reaction) and blank control (DW).

The analytical specificity of the *mcr-1-*MCDA-LFB assay was assessed with genomic DNA extracted from 16 *mcr-1*-producing strains and 35 non-*mcr-1* isolates. As shown in **Fig. 5**, all products derived from the strains carring *mcr-1* gene exhibited two red bands (TL and CL) in the LFB, but each sample from the *mcr-1-*negative organisms and blank control was only one red band. The results certified that the MCDA-LFB assay had a complete specificity for *mcr-1* detection.

**FIG 5.**
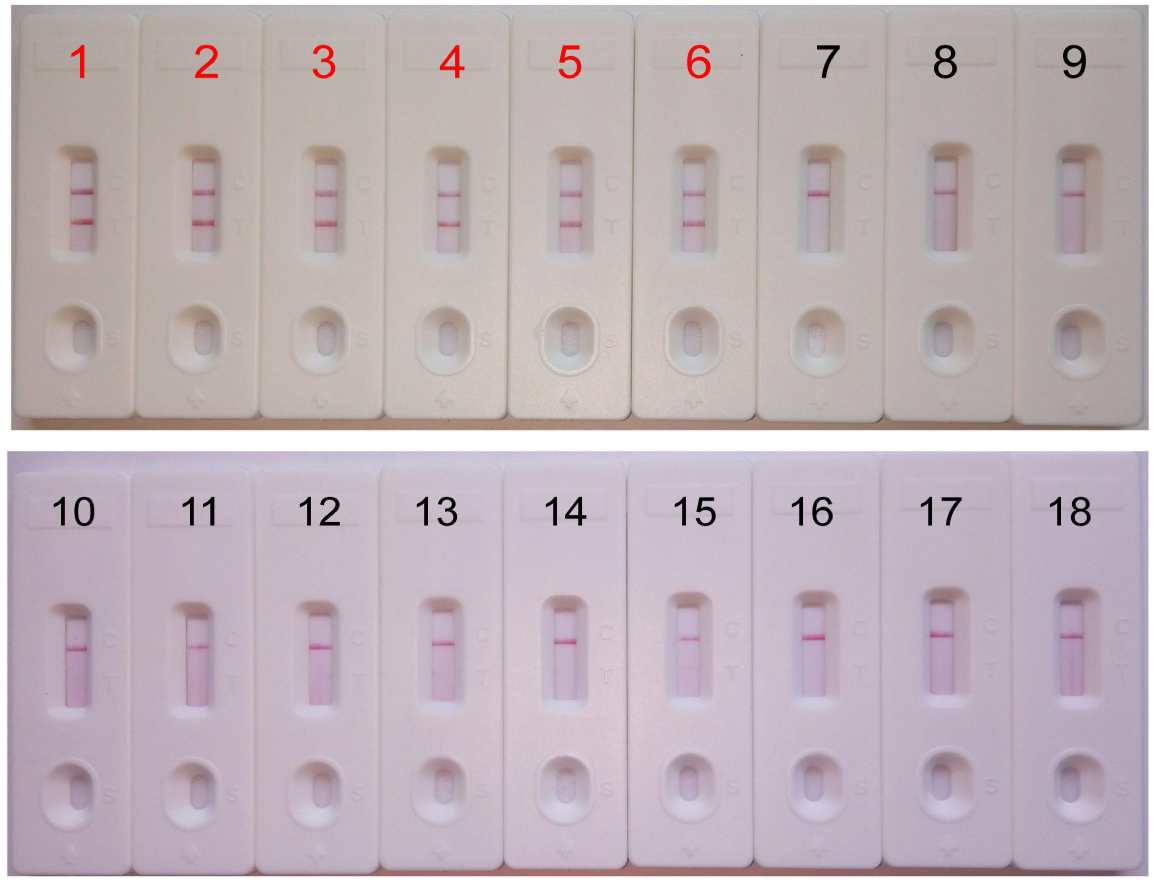
Specificity of the *mcr-1*-MCDA-LFB assay. Biosensor 1, *mcr-1*-positive *Escherichia fergusonii* ICDC-ZG2016M34-3. Biosensor 2, *mcr-1*-positive *Escherichia fergusonii* from ICDC. Biosensor 3, *mcr-1*-positive *Escherichia fergusonii* from WHCDC. Biosensor 4, *mcr-1*-producing *Escherichia coli* from ICDC. Biosensor 5, *mcr-1*-producing *Escherichia coli* from WHCDC. Biosensor 6, *mcr-1*-producing *Salmonella enteritidis* from ICDC. Biosensors 7-17, KPC-2-postive *Klebsiella pneumoniae,* KPC-2-postive *Pseudomonas aeruginosa,* NDM-1-postive *Escherichia coli,* NDM-1-postive *Enterobacter cloacae,* NDM-1-postive *Klebsiella pneumoniae,* NDM-5-postive *Escherichia coli,* IMP-4-postive *Escherichia coli, Acinetobacter baumannii* carring non*-mcr-1/*non*-*carbapenemase gene, *Pseudomonas aeruginosa* carring non*-mcr-1/*non carbapenemase gene, *mcr-1/*carbapenemase-negative *Serratia marcescens, mcr-1/*carbapenemase-negative *Escherichia coli,* all of the stains derived from WHCDC. Biosensor 18, blank control (DW).

### Optimization of the time for *mcr-1*-MCDA-LFB assay

To evaluate the optimum time, four reaction times (10 min to 40 min at 10 min intervals) were tested for the *mcr-1*-MCDA-LFB assay during the amplification stage. The *mcr-1*-producing *Escherichia fergusonii* (ICDC-ZG2016M34-3) plasmid DNA, 600 fg/μl (the LOD of the method), did not gain the positive results until the reaction had operated for 30 min (**Fig. 6**). Hence, the amplification time of 30 min at 63°C was considered as an optimal reaction condition for the current assay.

**FIG 6.**
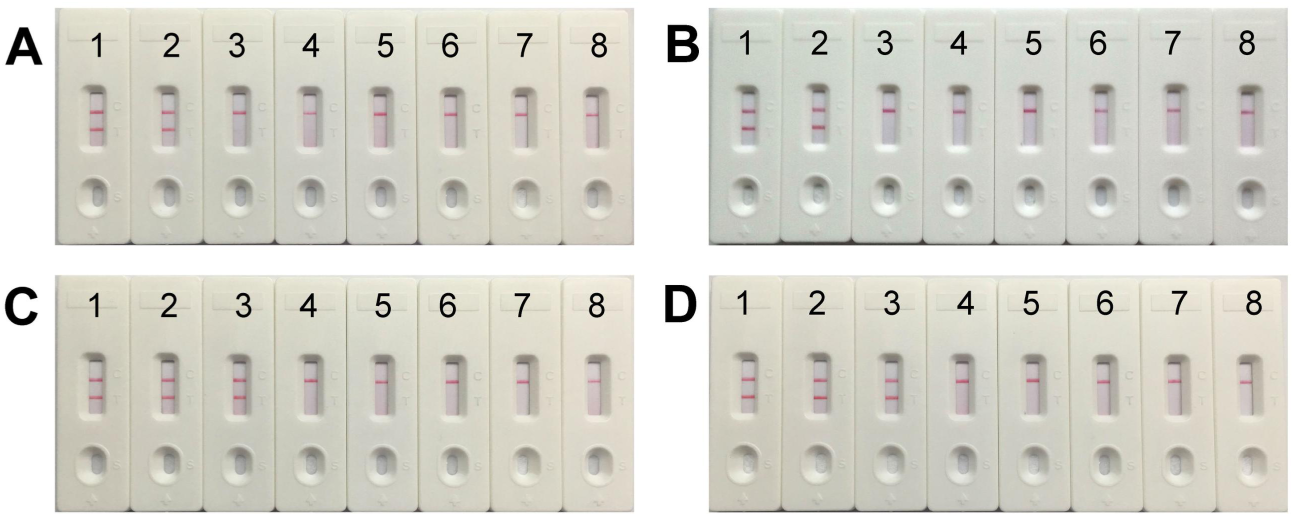
The optimal reaction time for *mcr-1*-MCDA-LFB assay. Four different amplification times, covering 10min (A), 20min (B), 30min (C) and 40min (D), were compared at 63°C. Biosensors 1-8 represented the plasmid DNA levels of 60 pg, 6 pg, 600 fg, 60 fg and 6 fg of *Escherichia fergusonii* ICDC-ZG2016M34-3, negative control (6 pg of NDM-1-positive *Escherichia coli* WHCDC-WH67 genomic DNA), negative control (6 pg of KPC-2-producing *Klebsiella pneumoniae* WHCDC-WH108 genomic DNA) and blank control (DW).

### Application of MCDA-LFB to *mcr-1*-spiked fecal samples

The LOD for strains expressing *mcr-1* in fecal samples was determined to assess the practical application of the established assay. The detected threshold of *mcr-1*-positive bacteria was approximately 4.5×10^3^ CFU/mL (~4.5CFU/reaction) in 0.2g faecal samples spiked with 100 μl of dilutions of strains (**Fig. 7**). The results of other subjects including the lower suspensions concentrations, negative control and blank control were negative. As the same to the aforementioned experiments, detection of the amplicons with three methods got an equal conclusion.

**FIG 7.**
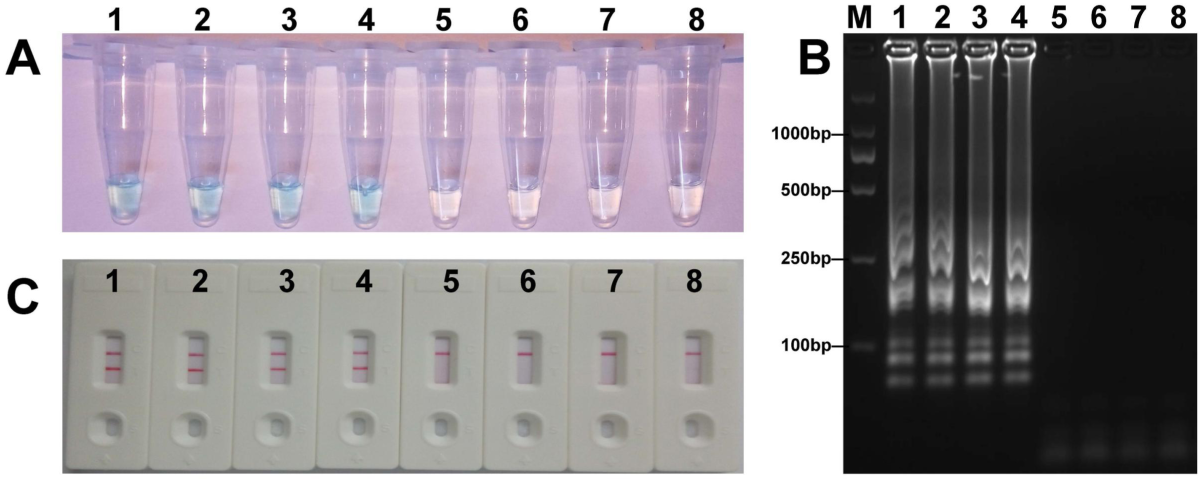
Sensitivity of *mcr-1-*MCDA-LFB assay in spiked feces samples. Three measurement techniques, including colorimetric indicators (A), gel electrophoresis (B) and LFB (C), were applied for testing MCDA products. Tubes (A)/lanes (B)/biosensors(C) 1-8 represented the DNA levels of 4.5 × 10^3^ CFU, 4.5 × 10^2^ CFU, 4.5 × 10^1^ CFU, 4.5 × 10^0^ CFU, 4.5 × 10^−1^ CFU and 4.5 × 10^−2^ CFU per reaction, negative control (non-spiked feces sample) and blank control (DW).

## Discussion

Polymyxins (polymyxin B and colistin) have nearly become last-resort drugs for treating the severe infections caused by multidrug-resistant or pan-resistant *Enterobacteriaceae* (6). The defensive line will be destroyed by the emergence of *mcr-1*-positive strains resisting to colistin. Moreover, the *mcr-1* gene mediated by plasmids or transposons can transfer in different species freely. Thus, the isolates carrying *mcr-1* gene will undoubtedly become a major issue for public health. Under the circumstances, a convenient and fast technique for detection of *mcr-1* in various samples is of great importance. Here, an approach was reported to detect target gene by multiple cross displacement amplification united with lateral flow biosensor (MCDA-LFB). In the MCDA-LFB assay, the high specificity was guaranteed, as a set of 10 primers was employed for specially amplifying the target sequence. The specificity of *mcr-1*-MCDA-LFB was successfully confirmed with the genomic templates extracted from *mcr-1*-producing strains and non-*mcr-1* organisms, and the results were positive for all *mcr-1*-positive isolates, but negative for non-*mcr-1* isolates and blank control. Therefore, the diagnostic test based on MCDA-LFB for the detection of *mcr-1* in bacteria identifies target gene with high selectivity.

The MCDA products can be analyzed with LFB, colorimetric indicator and agarose gel electrophoresis, respectively. LFB, as a detection technique by observing the number of red lines on sensor bar, is more objective than colorimetric indicator, which reports the result through color change. Maybe the latter is in trouble when the concentration of target gene is very low (24). Likewise, LFB is more rapid and convenient than gel electrophoresis, which requires the use of an additional operation procedure and complex equipment. Hence, LFB will be a better candidate for the results’ judge of MCDA products.

Besides specificity, the superb sensitivity is also very important for the newly established assay. The *mcr-1*-MCDA-LFB method sufficed to detect as little as 600 fg of *mcr-1*-positive plasmid DNA per microliter in pure culture and 4.5×10^3^ CFU/mL (~4.5 CFU/reaction) in fecal samples spiked with 100 μl of strains. This technique has the same sensitivity to *mcr-1*-LAMP described in previously report (25). Both isothermal amplification assays are 10 times more sensitive than conventional PCR (25). Although the Real-time PCR and MALDI-TOF MS-based method could also test the *mcr-1* gene with low limit of detection and less time, respectively (18, 26), they needed expensive apparatus and immaculately experimental condition that were not well equipped in resource-challenged fields, especially in the livestock industry where a large quantity of stains carrying *mcr* gene were identified (26).

The *mcr-1*-MCDA-LFB assay only required a constant reaction temperature at 63°C. The entire process of experiments, including sample processing (25 min), isothermal amplification (30 min) and detection (2min), could be accomplished less than 60 min. Herein, this assay economizes the test time and device, and is suitable for timely identification on the spot particularly.

In conclusion, we devised a reliable MCDA-LFB assay for the detection of *mcr-1* with simplicity, rapidity, and low-cost facility. The LOD of this assay was only 600 fg per reaction with pure culture, and the specificity was 100% according to the trial results. From the above, the *mcr-1*-MCDA-LFB assay built in this study will greatly improve the detection efficiency for the monitor of target gene in practical application.

## Acknowledgements

We acknowledge the financial supports (the research projects of clinical medicine in Wuhan, Grant Number:WG17Q02) from the Wuhan Municipal Health Commission, People’s Republic of China. We would like to thank Prof. Juan Li and Jie Che(Chinese Center for Disease Control and Prevention) for providing partial isolates. We are also grateful to Dr. Yi Wang (Beijing Children’s Hospital) for affording the technical guidance.

